# Differential susceptibility of human microglia HMC3 cells and brain microvascular endothelial HBEC-5i cells to Mayaro and Una virus infection

**DOI:** 10.1101/2023.09.02.556065

**Authors:** Dalkiria Campos, Madelaine Sugasti-Salazar, Paola Elaine Galán-Jurado, Patricia Valdés-Torres, Dalel Zegarra, José González-Santamaría

## Abstract

Mayaro (MAYV) and Una (UNAV) are emerging alphaviruses circulating in the Americas. Earlier reports have revealed that MAYV infects different human cell lines, including synovial and dermal fibroblasts, chondrocytes, osteoblasts, astrocytes and pericytes, as well as neural progenitor cells. In this study we evaluated the susceptibility of immortalized human microglia HMC3 cells and brain microvascular endothelial HBEC-5i cells to MAYV and UNAV infection. Cytopathic effects, cell viability, viral progeny yields, and the presence of E1 and nsP1 proteins in HMC3 and HBEC-5i cells infected with several MAYV or UNAV strains were assessed using an inverted microscope, MTT assay, plaque-forming assays, and immunofluorescence or Western blot, respectively. Finally, the expression of immune response genes was analyzed using qRT-PCR. MAYV and UNAV demonstrated strong cytopathic effects and significantly reduced cell viability in HMC3 cells. Moreover, the HMC3 cells were efficiently infected regardless of the virus strains tested, and E1 and nsP1 viral proteins were detected. In contrast, only MAYV appeared to infect HBEC-5i cells, and minimal effects on cell morphology or viability were observed. Furthermore, the MAYV titer and viral protein levels were substantially lower in the infected HBEC-5i cells when compared to those of the infected microglia cells. Finally, unlike UNAV, MAYV elicited a strong expression of specific interferon-stimulated genes in microglia cells, along with pro-inflammatory cytokines implicated in the immune response. Collectively, these findings demonstrate that MAYV and UNAV are capable of infecting relevant human brain cells.

## 1. INTRODUCTION

Mayaro (MAYV) and Una (UNAV) viruses are emerging and neglected New World alphaviruses within the *Togaviridae* family [1]. MAYV is the etiologic agent responsible for Mayaro fever, a disease with unspecific symptoms, including fever, headache, rash, leukopenia, diarrhea, myalgia, retroorbital pain and some cases, a severe polyarthralgia that can last for months to years [1,2]. In contrast, there is only serologic evidence of UNAV infection in human or non-human primates, and the symptoms associated with this infection are unknown [3-5]. MAYV is actively circulating in different countries of Central and South America, mostly through an enzootic cycle in which the primary vectors are sylvatic mosquitoes *Haemagogus janthinomys* and the main hosts are non-human primates [6,7]. Nevertheless, laboratory and field studies suggest that urban vectors, such as *Aedes aegypti* or *Aedes albopictus*, have the capacity to transmit this virus [8,9]. On the other hand, several strains of UNAV have been isolated from *Psorophora ferox* and *Psorophora albipes* mosquitos in different Latin American countries [10-12]. Although MAYV and UNAV infection represent a potential public health threat, both viruses are poorly characterized pathogens and obtaining new information regarding viral pathogenesis and cell tropism remains fundamental.

In this context, Cavalheiro and colleagues demonstrated that MAYV were able to infect mouse primary macrophages and mouse macrophage-derived cell lines in a time-dependent manner [13]. In these cells, MAYV infection induced the expression of tumor necrosis factor alpha (TNF-α) and the production of reactive oxygen species (ROS), contributing to cell death [13]. In another study using human hepatic HepG2 cell lines, the authors also found that MAYV infection promoted ROS production, and consequently, provoked oxidative stress in these cells [14]. It has been reported that MAYV induces a persistent polyarthralgia in some infected patients [2]. Thus, Bengue and coworkers evaluated the susceptibility of human chondrocytes, fibroblast-like synoviocytes and osteoblasts, the main cell types implicated in osteoarthritis, to MAYV. Their results revealed that all these cell lines could be infected with MAYV [15]. Moreover, several arthritis-related genes were upregulated in infected human chondrocytes, such as matrix metalloproteinases (MMP) including MMP1, MMP7, MMP8, MMP10, MMP13, MMP14 and MMP15 [15]. Our group found that primary human dermal fibroblasts are susceptible to MAYV infection and elicit a strong expression of interferon-stimulated genes (ISGs), interferon, and cytokines, promoting an antiviral state [16]. To the best of our knowledge, there are no previous studies analyzing the cell tropism of UNAV.

Although MAYV infection has not been associated with neurological sequelae, recent reports indicate that MAYV efficiently infect human neural progenitor cells, pericytes and astrocytes *in vitro* [17,18]. Moreover, in astrocytes MAYV infection induces a strong antiviral response and the production of inflammatory mediators, which could contribute to neuroinflammation [17]. Nevertheless, whether MAYV or UNAV can infect other key brain cells remains unexplored. Microglia are immune cells in the central nervous system (CNS) that play an important role during viral infection and in inflammatory diseases [19]. In addition, brain microvascular endothelial cells are fundamental elements in the blood-brain barrier, a highly selective and semipermeable barrier that prevents the free passage of solutes or blood into the CNS [20]. In the present study we evaluated the susceptibility of human microglia HMC3 cells and brain microvascular endothelial HBEC-5i cells to MAYV and UNAV infection.

## 2. MATERIAL AND METHODS

### 2.1 Cell lines, virus strains and reagents

Human microglia HMC3 cells (CRL-3304), human brain microvascular endothelial HBEC-5i cells (CRL-3245) and Vero-E6 cells (CRL-1586) were obtained from American Type Culture Collection (ATCC, Manassas, VA, USA) and grown in Eagle’s Minimum Essential Medium (EMEM), Dulbecco’s Modified Eagle’s Medium (DMEM) / Kaighn’s Modification of Ham’s F-12 Medium (F-12K), and Minimal Essential Medium (MEM), respectively. All media were supplemented with 10% fetal bovine serum (FBS), a 1% penicillin-streptomycin antibiotic solution, and 2mM of L-Glutamine (all reagents were obtained from Gibco, Waltham, MA, USA). For the endothelial cells, the medium was supplemented with 100 µg/ml of Endothelial Cell Growth Supplement (Millipore, Temecula, CA, USA). Cell lines were incubated at 37 °C under a 5% CO2 atmosphere. The Mayaro (MAYV, AVR 0565, Peru; MAYV, Guyane, French Gianna; and MAYV, TRVL 4675, Trinidad and Tobago) and Una (UNAV, BT 1495-3, Panama; UNAV, 788382, Trinidad and Tobago; and UNAV, CoAr 2518, Colombia) strains used in this work were obtained from the World Reference Center for Emerging Viruses and Arboviruses (WRCEVA) at University of Texas Medical Branch (UTMB, USA) and kindly provided by Dr. Scott Weaver. All virus strains were produced in Vero-E6 cells and then titrated, aliquoted and stored as previously reported [21].

### 2.2 Cytopathic effect and cell viability assays

HMC3 and HBEC-5i cells were grown in 12-well or 96-well plates and then infected with MAYV or UNAV at a multiplicity of infection (MOI) of 1 for a period of 24 to 72 h. After that, images were captured with an inverted microscope to analyze cell morphology. For the cell viability experiments, we used the MTT method as previously described [21]. After 24, 48 or 72 h of infection, 5 mg/ml of 3-(4,5-Dimethyl-2-thiazolyl)-2,5-diphenyltetrazolium bromide (MTT, Sigma-Aldrich, St. Louis, MI, USA) solution was added to the cells and incubated for an additional 4 h. Formazan crystals were dissolved in a solution of 4 mM HCl and 10% Triton X-100 in isopropanol, and absorbance was determined at 570 nm using a microplate reader spectrophotometer (BioTeK, Winooski, VT, USA). Results are reported as the percentage of viable cells relative to untreated control cells.

### 2.3 Virus titration using plaque-forming assay

Virus progeny production in cell supernatants was determined using plaque-forming assays as previously performed [21]. For this end, 10-fold serial dilutions of MAYV- or UNAV-infected samples were used to infect confluent Vero-E6 cells grown in 6-well plates. Next, the inoculum was eliminated, and the cells were overlaid with a solution of 1% agar in MEM supplemented with 2% FBS; then, the cells were incubated at 37 °C for 3 days. Subsequently, the agar was removed, and the cells were fixed with 4% formaldehyde solution in PBS and stained with 2% crystal violet dissolved in 30% methanol solution. Lastly, the number of plaques were counted, and the viral titers were reported as plaque-forming units per milliliter (PFU/ml).

### 2.4 Immunofluorescence assay

HCM3 or HBEC-5i cells grown on glass coverslips in 24-well plates were infected with MAYV or UNAV at an MOI of 1 for 48 h. Next, the cells were fixed, permeabilized, and blocked as previously reported [21]. After that, the cells were stained overnight at 4 °C with E1 or nsP1 primary rabbit antibodies (both antibodies previously validated in our laboratory) [22], followed by Alexa Flour 568 goat anti-rabbit secondary antibody (Invitrogen, Carlsbad, CA, USA). Lastly, coverslips were mounted on slides with Prolong Diamond Antifade Mountant with DAPI (Invitrogen, Carlsbad, CA, USA), and images were obtained with an FV1000 confocal microscope (Olympus, Lombard, IL, USA). The images were analyzed with ImageJ software (National Institute of Health, Bethesda, MD, USA).

### 2.5 Western blot assay

Viral protein levels were evaluated by Western blot as previously described [21]. Protein extracts were obtained from MAYV- or UNAV-infected HMC3 or HBEC-5i cells in Laemmli buffer with 10% dithiothreitol (Bio-Rad, Hercules, CA, USA). Proteins were fractionated in SDS-PAGE, transferred to nitrocellulose membranes, and blocked with a solution of 5% non-fat milk in T-TBS buffer for 30 min. After, the membranes were incubated with the following primary antibodies: rabbit polyclonal anti-E1, rabbit polyclonal anti-nsP1, and mouse monoclonal anti-β-actin (Cat # VMA00048, Bio-Rad, Hercules, CA, USA). Next, the membranes were washed three times with T-TBS buffer and incubated with HRP-conjugated goat anti-rabbit (Cat. # 926-80011) or goat anti-mouse (Cat. # 926-80010) secondary antibodies (LI-COR, Lincoln, NE, USA) for 1 h at room temperature. Finally, the membranes were incubated with SignalFireTM ECL Reagent (Cell Signaling Technology, Danvers, MA, USA) for 5 min, and the chemiluminescent signal was detected with a C-Digit scanner (Li-COR, Lincoln, NE, USA).

### 2.6 Analysis of mRNA expression using qRT-PCR assay

Total RNA from mock- or infected-HMC3 cells were obtained using an RNeasy Kit (Qiagen, Valencia, CA, USA) following the manufacturer’s instructions. cDNA was synthesized from 1 µg of RNA using a High-Capacity cDNA Reverse Transcription Kit, and quantitative RT-PCR was performed using Power SYBR Green PCR Master Mix in a QuantStudioTM 5 thermocycler (Applied Biosystems, Foster City, CA, USA). The following immune response genes were detected: MDA5, OAS2, MxA, ISG15, AIM2, RIG-I, IFN-α, IFN-β, TNF-α, IL-6, IL-1β, IL-8, CCL5, IRF-3, IRF-7, TLR3, and TLR7. All genes were previously validated [23-28] and are listed in Table S1 (Supplementary Material). Relative mRNA expression was determined using the β-actin gene for normalization according to the ΔΔ CT method [29].

### 2.7 Data analysis

All data were analyzed with the Mann & Whitney test. Data analysis was performed, and graphics were created with GraphPad Prism software version 10.0.0 for Mac. All experiments were performed at least 2 or 3 times with 3 replicates. For each assay, the mean and standard deviation are shown. A p value < 0.05 was considered statistically significant.

## 3. RESULTS

### 3.1. Mayaro and Una viruses promote strong cytopathic effects and decrease cell viability in human microglial HMC3 cells but have little effect on brain microvascular endothelial HBEC-5i cells

Recently it was reported that MAYV efficiently infects different human brain cells, including neural progenitor cells, pericytes, and astrocytes [17]. To explore whether MAYV and its closest-related alphavirus, UNAV, have the capacity to infect other human brain cells, we evaluated microglia HMC3 cells and brain microvascular endothelial HBEC-5i cells, both of which are immortalized cells that preserve many of the characteristics of the primary cells from which they were derived. Previous studies have revealed that MAYV, like other alphaviruses, is capable of inducing a potent cytopathic effect in different cell lines, including Vero cells and primary human dermal fibroblasts [16,30]. Thus, we infected HMC3 and HBEC-5i cells with MAYV or UNAV at a multiplicity of infection (MOI) of 1, and at different times points infection, we analyzed the cell morphology using an inverted microscope. In this experiment we found that both MAYV and UNAV promoted substantial cytopathic effects in HMC3 cells in a time-dependent manner (Figure 1A). In contrast, we observed minor cytopathic effects in HBEC-5i cells infected with MAYV, and these cells appeared to be unaffected by UNAV infection (Figure 1A). To evaluate the impact of the MAYV or UNAV infection on the cellular integrity of HMC3 and HBEC-5i cells, we performed an infection kinetic experiment; at specific time points, cell viability was measured using the MTT method. As shown in the Figure 1, MAYV and UNAV infection provoked a significant reduction in cell viability among the HMC3 cells at 72 hours post infection (hpi) (Figure 1B). However, we observed a less pronounced decrease in cell viability among MAYV-infected HBEC-5i cells (Figure 1C). Together, these results indicate that MAYV and UNAV infection promote extensive cell damage in human microglia HMC3 cells.

**Figure 1.**
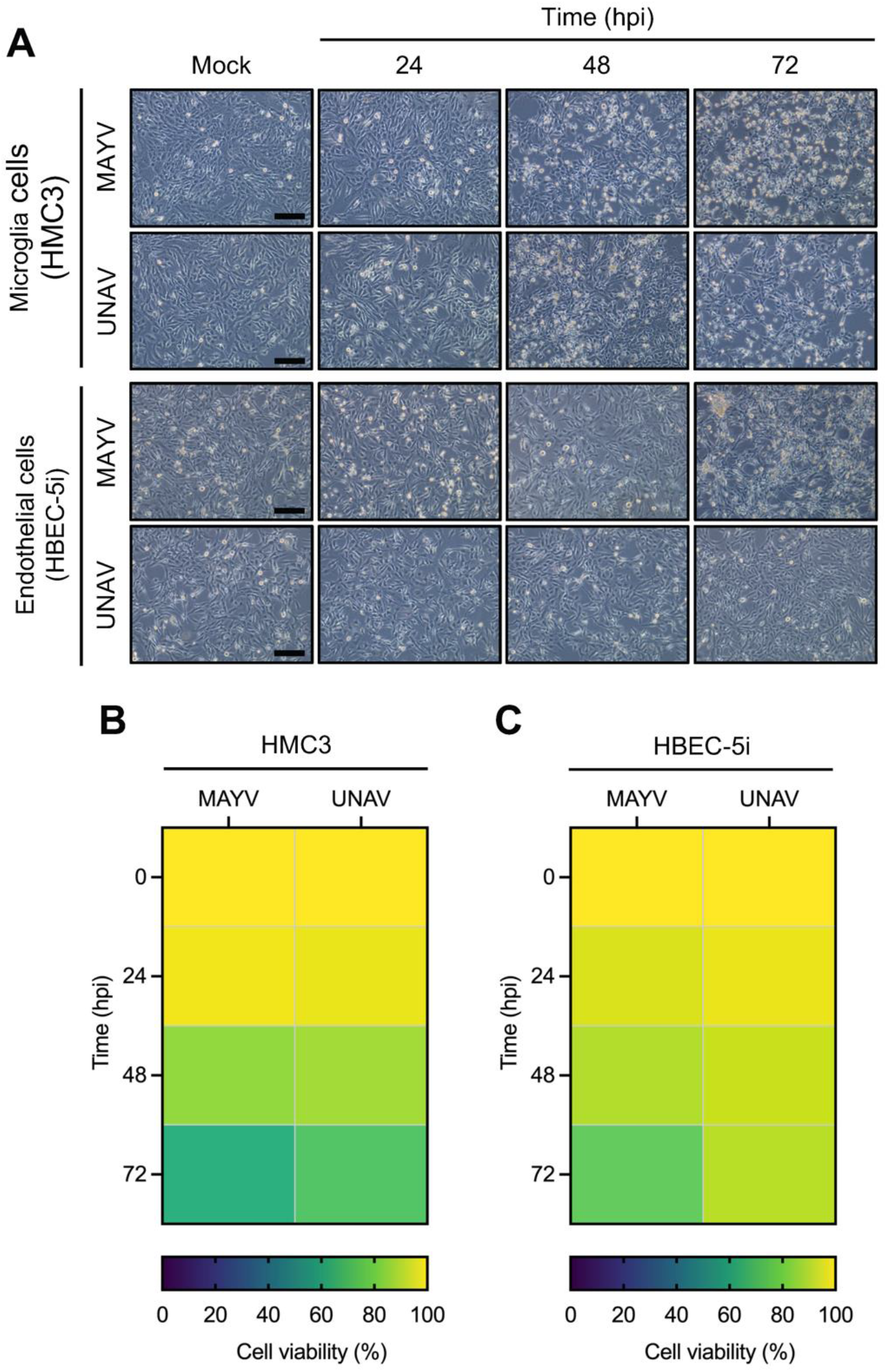
Mayaro and Una viruses promote cytopathic effects in and reduce cell viability of human microglia HMC3 cells. (**A**) Human microglia HMC3 cells or brain microvascular endothelial HBEC-5i cells were infected with MAYV (AVR 0565 strain) and UNAV (BT 1495-3 strain) at an MOI of 1. At the indicated times, cytopathic effects were evaluated using an inverted microscope. A representative image of at least 10 fields is shown. Scale bar: 100 µm. HMC3 (**B**) and HBEC-5i (**C**) cells were infected with MAYV or UNAV as described above. At different time points, cell viability was assessed using the MTT method; the results are presented as a heat map showing the percentage of viable cells.

### 3.2. Mayaro and Una viruses efficiently replicate in human microglia HMC3 cells in a time- and MOI-dependent manner

To evaluate whether MAYV and UNAV can replicate in HMC3 or HBEC-5i cells, we performed an infection kinetic experiment in which both cell lines were infected with MAYV or UNAV at a MOI of 1 or 10; at different time points, the viral progeny yields in cell supernatants were assessed using plaque-forming assays. In this experiment, we found higher viral titers in the supernatants of MAYV- and UNAV-infected microglia HMC3 cells (Figure 2A, B), and this effect was time- and MOI-dependent. On the contrary, while MAYV demonstrated efficacy in infecting brain microvascular endothelial HBEC-5i cells, although at a low level, UNAV did not effectively infect these cells (Figure 2C, D).

**Figure 2.**
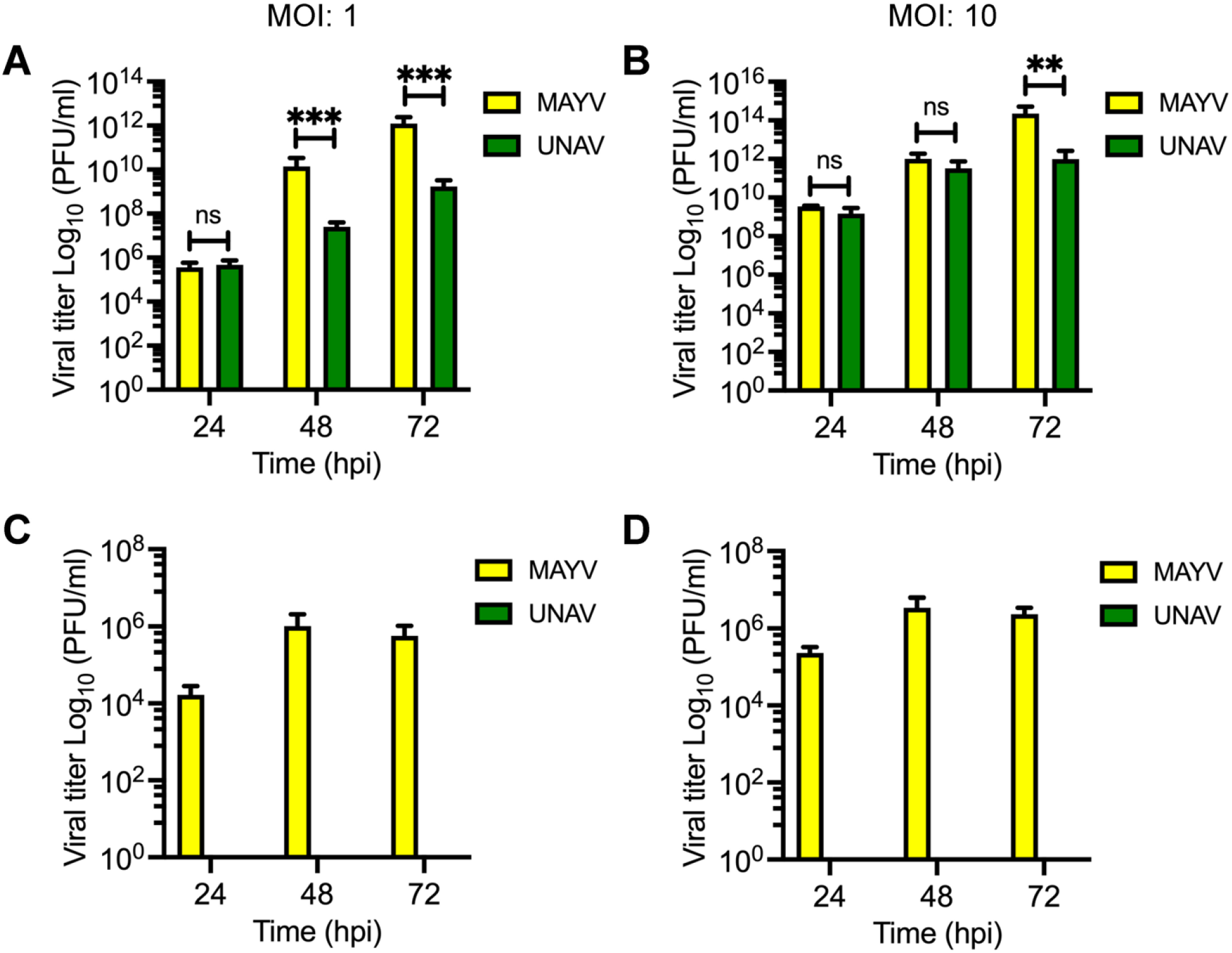
Mayaro and Una viruses infect human microglia HMC3 cells. HMC3 (**A, B**) or HBEC-5i (**C, D**) cells were infected with MAYV (AVR 0565 strain) or UNAV (BT 1495-3) at an MOI of 1 or 10 for 1h. Next, the inoculum was removed, and the cells were incubated for the indicated times. Viral progeny production in cell supernatants was quantified using plaque-forming assays. Viral titers are reported as plaque-forming units per milliliter (PFU/ml). The mean and standard deviation of three independent experiments in triplicate are shown, and the data were analyzed using the Mann & Whitney test. Statistically significant differences are denoted as follows: ** p < 0.01; *** p < 0.001; ns: non-significant.

To verify these previous findings, we decided to analyze the expression of the E1 and nsP1 viral proteins in HMC3 or HBEC-5i cells. We performed an immunofluorescence assay in both cell lines infected with MAYV or UNAV. In this experiment, we observed strong expression of both viral proteins in MAYV- or UNAV-infected microglia HMC3 cells at 48 hpi (Figure 3A). In the case of the brain endothelial HBEC-5i cells, we noted a low expression of both proteins in MAYV-infected cells, but not expression was observed in UNAV-infected cells. To further corroborate the results, we evaluated the levels of both viral proteins in MAYV- or UNAV-infected HMC3 or HBEC-5i cells using Western blot. The infection kinetic assay revealed that there were high levels of the E1 and nsP1 proteins for both viruses at 48 hpi in microglia HMC3 cells (Figure 3B-C). In contrast, we did not detect any viral proteins in MAYV- or UNAV-infected HBEC-5i cells. Taken together, these results demonstrate that MAYV and UNAV competently infect human microglia HMC3 cells.

**Figure 3.**
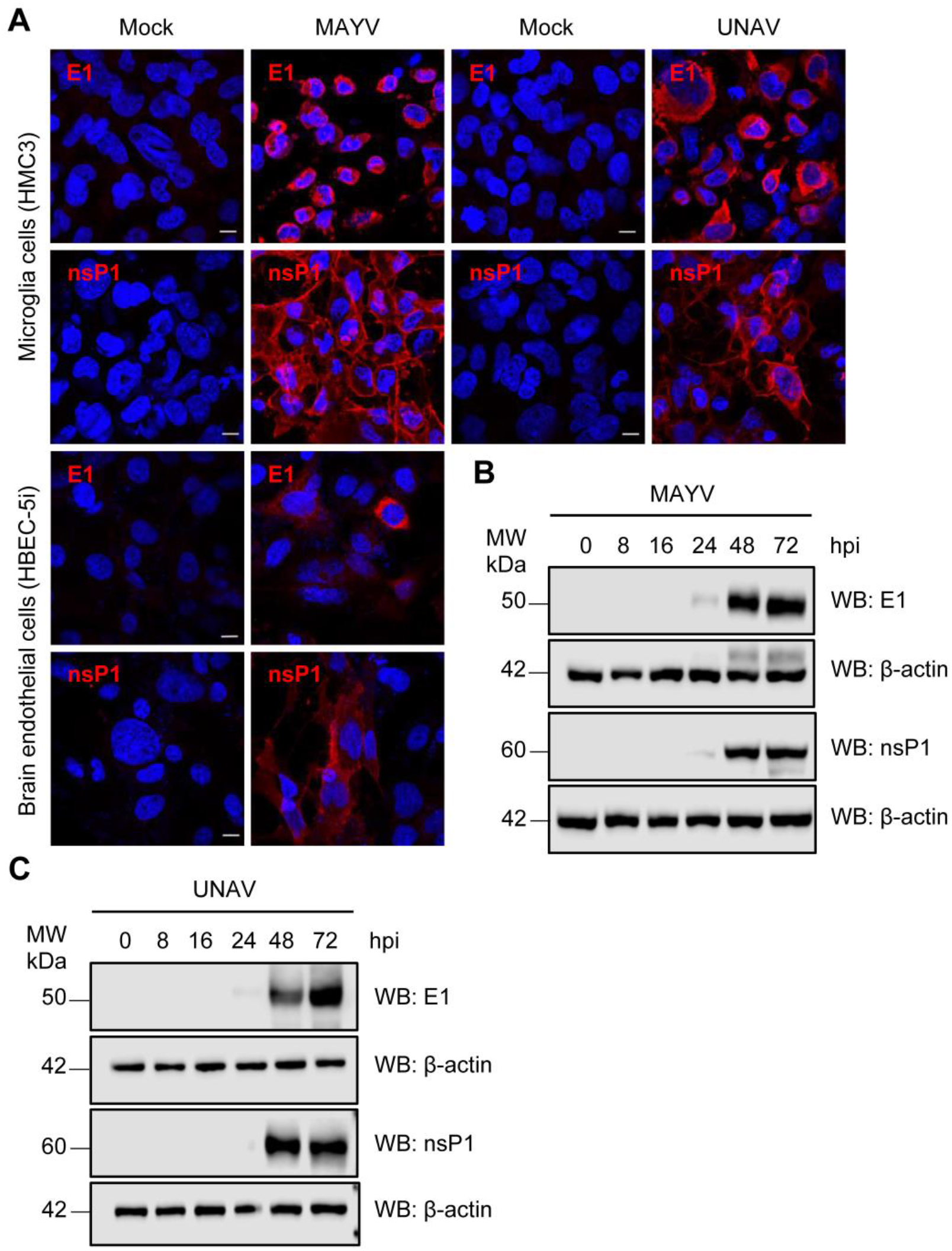
Detection of viral E1 and nsP1 proteins in MAYV- or UNAV-infected HMC3 and HBEC-5i cells. (**A**) HMC3 and HBEC-5i cells were infected with MAYV (AVR 0565 strain) or UNAV (BT 1495-3 strain) as previously performed. Following 48h of infection, the viral E1 and nsP1 proteins were assessed by immunofluorescence. A representative image of at least 10 fields is shown. Kinetic infection assay in HMC3 cells infected with MAYV (**B**) or UNAV (**C**) showing the E1 and nsP1 protein levels using Western blot. A representative blot of three independent experiments is shown. β-actin antibody was used as a loading control. WB: Western blot; MW: molecular weight; kDa: kilodaltons.

### 3.3 All tested Mayaro and Una virus strains infect human microglia HMC3 cells

To validate our preceding results, we decided to assess the capacity of other MAYV and UNAV strains to infect HMC3 and HBEC-5i cells. We infected both cell lines with the MAYV Guyane and TRVL 4675 or UNAV 788382 and CoAr 2518 strains, all isolated from different Latin American countries. As we observed in our previous experiments, all the MAYV or UNAV strains tested were able to efficiently infect human microglia HMC3 cells in a time-dependent manner (Figure 4A-B). Again, only the MAYV strains infected the brain endothelial HBEC-5i cells, and with less efficiency (Figure 4C-D). These results confirm that MAYV and UNAV competently infect human microglia HMC3 cells.

**Figure 4.**
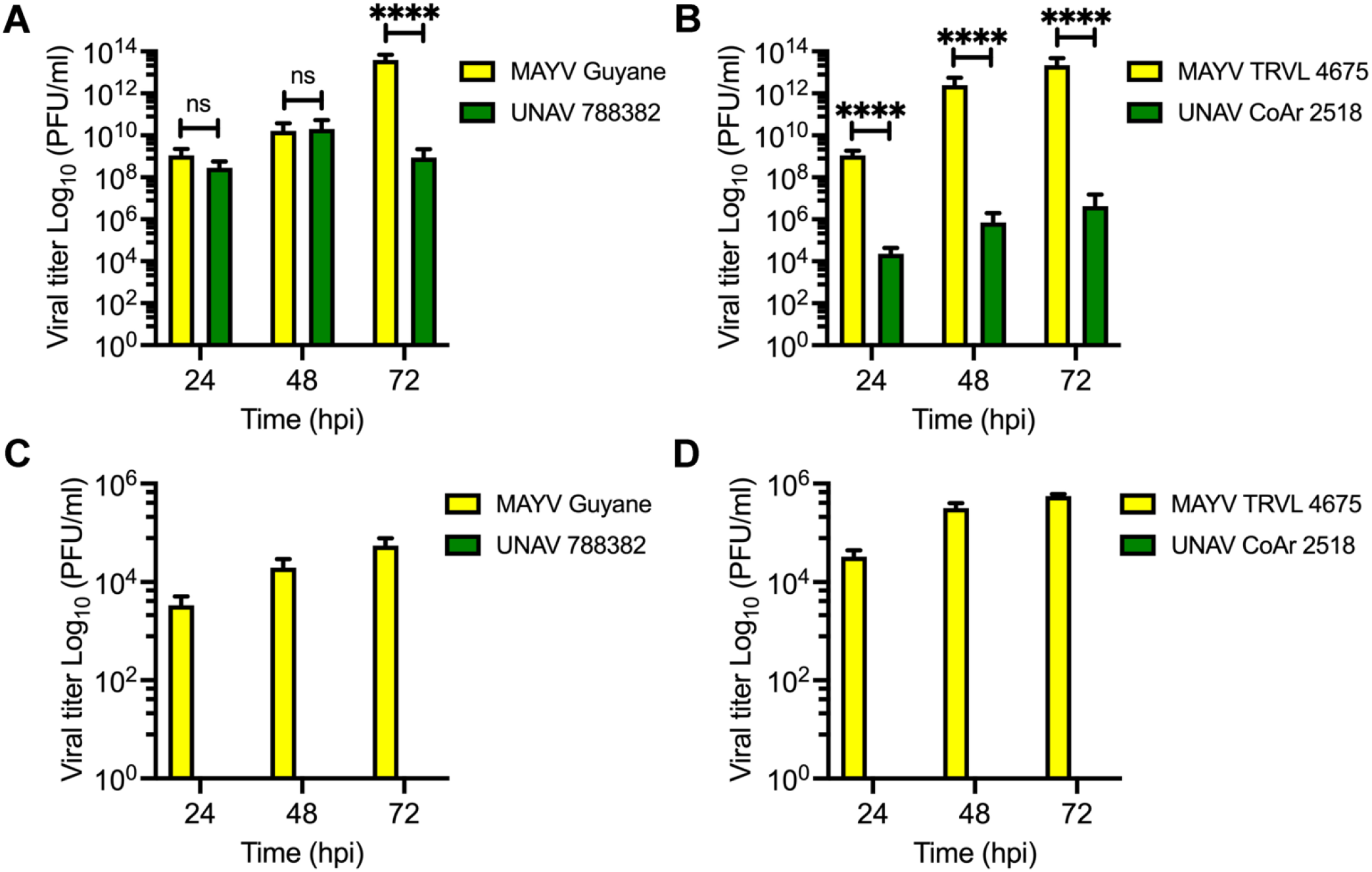
Different strains of Mayaro and Una viruses efficiently infect human microglia HMC3 cells. HMC3 (**A, B**) or HBEC-5i (**C, D**) cells were infected with MAYV Guyane and TRVL 4675 or UNAV 788382 and CoAr 2518 strains; at the indicated times points, viral progeny yields in cell supernatants were assessed using plaque-forming assays. Viral titers are reported as plaque-forming units per milliliter (PFU/ml). The mean and standard deviation of three independent experiments in triplicate are shown, and the data were analyzed using the Mann & Whitney test. Statistically significant differences are denoted as follows: **** p < 0.0001; ns: non-significant.

### 3.4. Mayaro virus but not Una virus potently induces the expression of immune response genes in human microglia HMC3 cells

Given that human microglia HMC3 cells were high susceptible to MAYV and UNAV infection, we decided to focus on this cell line for a more detailed characterization of the infection process with these viruses. We analyzed the expression of immune response genes, including interferon, interferon-stimulated genes, cytokines, chemokines, and transcription factors in MAYV- or UNAV-infected HMC3 cells. As shown in Figure 5, we found that MAYV infection strongly triggers the expression of IFN-α and IFN-β, as well as the interferon-stimulated genes MDA5, OAS2, MxA, AIM2, and RIG-I (Figure 5). In addition, we observed an increase in expression of the cytokines TNF-α, IL-6, and IL-8, the transcription factors IRF-3 and IRF-7, and the toll-like receptors TLR3 and TLR7 genes (Figure 5). In the case of UNAV-infected HMC3 cells, we noted a less pronounced rise in the expression of MDA5, OAS2, MxA, IFN-α, IFN-β, TNF-α, IL-6, IL-8, CCL5, TLR3, and TLR7 genes (Figure 5). These findings suggest that MAYV induces a potent antiviral state in HMC3 cells.

**Figure 5.**
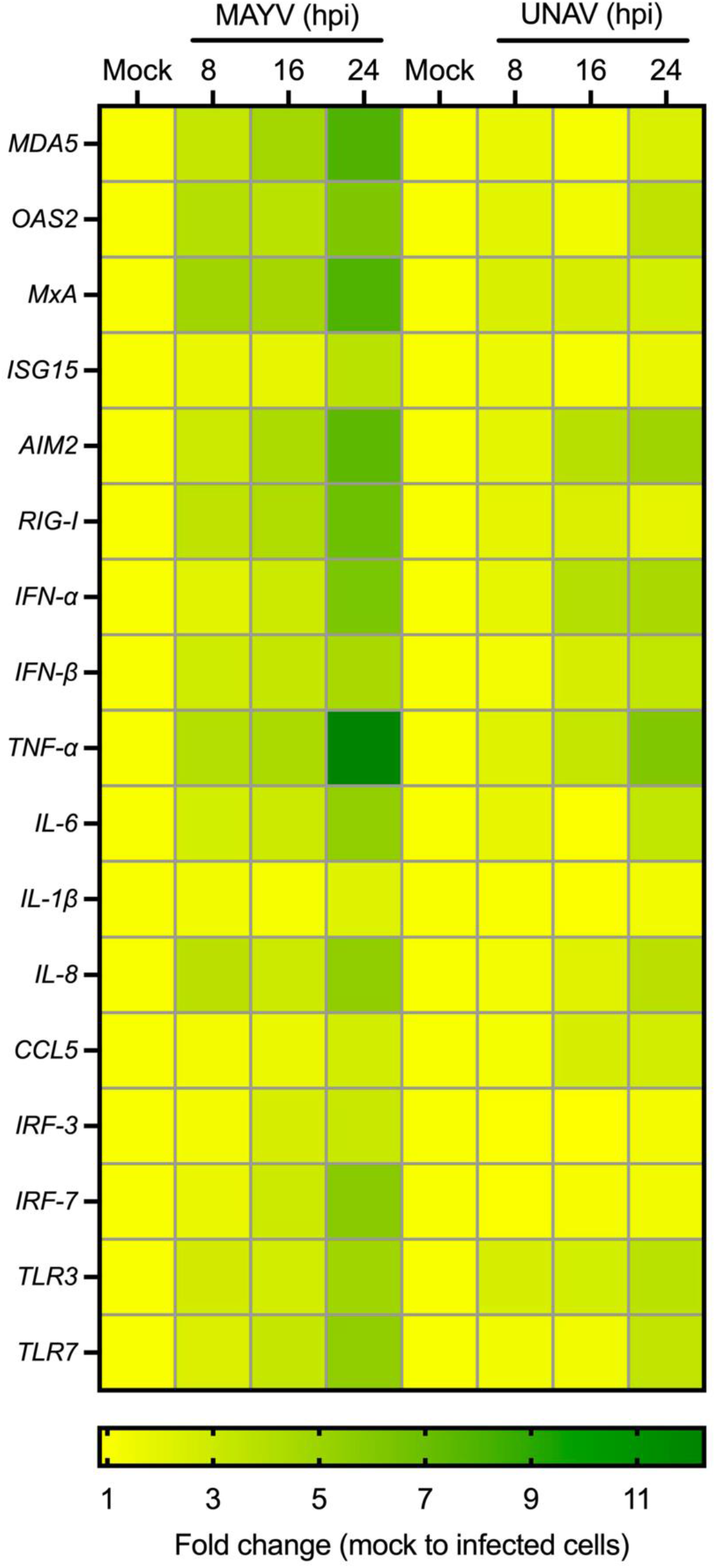
MAYV induces the expression of immune response genes in human microglia HMC3 cells. HMC3 cells were infected with MAYV (AVR 0565 strain) or UNAV (BT 1495-3) at an MOI of 1. At specific time points, expression of the indicated immune response genes was evaluated using qRT-PCR. The heat map expresses the fold change of specific genes when comparing mock control cells vs. MAYV- or UNAV-infected cells.

## 4. DISCUSION

MAYV and UNAV are emerging and neglected alphaviruses. The poor characterization of these viruses’ pathogenesis leaves a knowledge gap regarding their cell tropism. However, several *in vitro* studies demonstrate that MAYV can infect different human cell lines, among them, chondrocytes, fibroblast-like synoviocytes, osteoblasts, and dermal fibroblasts [15,16]. Moreover, it was recently reported that MAYV is able to infect several types of human brain cells, including pericytes, astrocytes, and progenitor neural cells [17]. Conversely, to the our best knowledge, there is not information available regarding UNAV’s cell tropism.

In the present study we evaluated the susceptibility of immortalized human microglia HMC3 cells and brain microvascular endothelial HBEC-5i cells to MAYV and UNAV infection. Our results indicate that MAYV as well as UNAV promoted substantial cytopathic effects in microglia HMC3 cells in a time-dependent manner. Also, we observed a significant reduction in cell viability among the HMC3 cells infected with these viruses. In contrast, only MAYV appeared to have an effect on brain microvascular endothelial HBEC-5i cells, and this effect was less pronounced.

To verify if HMC3 or HBEC-5i cells might be susceptible to MAYV and UNAV infection, we performed an infection kinetic experiment; at specific time points, we quantified viral progeny production in the supernatants of both cell lines. In these assays, we found that microglia HMC3 cells were vulnerable to infection with all MAYV and UNAV strains tested. In addition, we detected a strong expression of the viral E1 and nsP1 proteins in MAYV- or UNAV-infected HMC3 cells, corroborating our previous findings. However, only MAYV strains were able to infect brain microvascular endothelial HBEC-5i cells, and with a very limited capacity. A previous study demonstrated that Chikungunya virus (CHIKV), another arthritogenic alphavirus closely related to MAYV and UNAV, was able to infect microglia HMC3 cells [31].

To further characterize the effect of MAYV and UNAV infection in microglia HMC3 cells, we assessed the expression of immune response genes using qRT-PCR. Our data indicate that unlike UNAV, MAYV infection potently induced the expression of interferon, interferon-stimulated genes, and pro-inflammatory cytokines, promoting a general antiviral state. Similar results have been observed in astrocytes and human dermal fibroblasts infected with MAYV [16,17].

Although MAYV has not been associated with neurological sequelae clinically, our findings indicate that MAYV is able to infect, at least *in vitro*, other vital brain cells, such as microglia and endothelial cells. It is important to highlight that CHIKV has been implicated with neurological complications in human infection, including encephalopathy, encephalitis, encephalomyelopathy, myeloneuropathy, acute disseminate encephalomyelitis, and Guillain-Barré syndrome, among others [32]. Thus, studying the capacity of MAYV or UNAV to infect relevant human brain cells and their potential ability to produce neurological complications is of particular interest. In this context, it would be valuable to know if these viruses can infect neurons, given that at least MAYV has the capacity to infect brain endothelial cells. However, animal models of infection with MAYV or UNAV are required to validate the possible roles of these viruses in neuropathogenesis.

## Author Contributions

Conceptualization, D.C. and J.G.-S.; methodology, D.C., M.S.-S., P.V-T., P.E.G.-J., D.Z., and J.G.-S.; validation, D.C. and J.G.-S.; formal analysis, D.C. and J.G.-S.; investigation, D.C., M.S.-S., P.V.-T., P.E.G.-J., D.Z., and J.G.-S.; resources, J.G.-S.; writing—original draft preparation, J.G.-S.; writing—review and editing, D.C., M.S.-S., P.V.-T., P.E.G.-J., D.Z., and J.G.S; supervision, D.C. and J.G.-S.; project administration, J.G.-S.; funding acquisition, J.G.-S. All authors have read and agreed to the published version of the manuscript.

## Acknowledgments

We thank Scott Weaver (WRCEVA, UTMB, USA) for providing the Mayaro and Una virus strains. We also give thanks to Rodolfo Contreras and Nicanor Obaldía for their support with laboratory and equipment facilities. Finally, we express our gratitude to Jorge Ceballos and the Smithsonian Tropical Research Institute for their assistance and granting access to the confocal microscope.

## SUPPLEMENTARY MATERIAL

**Table S1.**
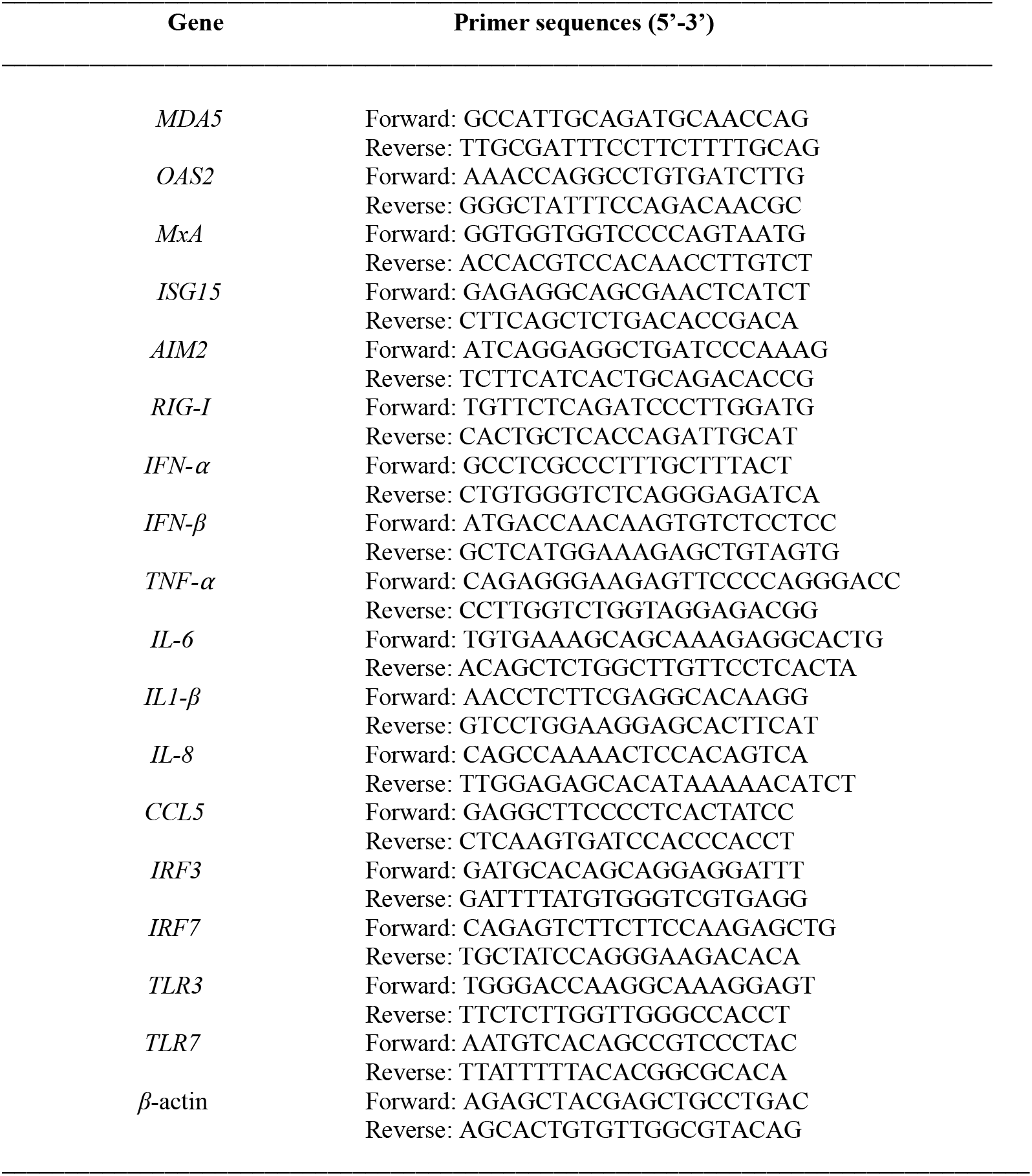
List of primers used in this study.

